# Low contribution of rare species to resilience and adaptive capacity in novel spatial regimes arising from biome shifts caused by global change

**DOI:** 10.1101/2020.01.29.924639

**Authors:** David G. Angeler, Caleb P. Roberts, Dirac Twidwell, Craig R. Allen

**Affiliations:** Department of Aquatic Sciences and Assessment, Swedish University of Agricultural Sciences, Box 7050, 750 07 Uppsala, Sweden; School of Natural Resources, University of Nebraska—Lincoln, Lincoln, NE 68583, USA; Department of Agronomy & Horticulture, University of Nebraska, Keim Hall, Lincoln, NE 68583, USA; Center for Resilience in Agricultural Working Lands, University of Nebraska, Lincoln, NE 68583, USA

**Keywords:** Great Plains, regime shifts, biome change, landscape ecology, biogeography, conservation, spatial regimes, resilience

## Abstract

Human activity causes biome shifts that alter biodiversity and spatial resilience patterns, ultimately challenging conservation. Rare species, often considered vulnerable to change and endangered, can be a critical element of resilience by providing adaptive capacity in response to disturbances. However, little is known about changes in rarity and dominance patterns of communities once a biome transitions into a novel spatial regime, and how this affects conservation. We used time series modeling to identify species rarity and dominance patterns in an expanding terrestrial (southern) spatial regime in the North American Great Plains and another (northern) regime that will become encroached by the southern regime in the near future. In this approach, presumably rare and abundant species show stochastic and deterministic dynamics, respectively. We specifically assessed how stochastic species of the northern spatial regime influence the resilience and adaptive capacity of a novel spatial regime once it has been encroached by the southern regime by either becoming deterministic or staying stochastic. Using 47 years (1968 – 2014) of breeding bird survey data and a space-for-time substitution, we found half of the stochastic species from the northern regime to be either deterministic or stochastic in the southern regime. However, the overall contribution of these species to the community of the southern regime was low, manifested in marginal contributions to resilience and adaptive capacity of this regime. Also, none of these species were of conservation concern, suggesting limited need for revised species conservation action in the novel spatial regime. From a systemic perspective our result suggest that while stochastic species can potentially compensate for the loss of dominant species after disturbances and maintain the system in its current regime, they may only marginally contribute to resilience and adaptive capacity in a new spatial regime after fundamental ecological changes have occurred.

## Introduction

With ongoing environmental change in the Anthropocene, ecosystems are changing rapidly at local, regional and global scales (Vellend et al. 2017). For example, multiple global change drivers (e.g. climate, species invasions, agriculture) within the Great Plains of North America affect ecosystems in a south-to-north pattern (Roberts et al. 2019). In the Great Plains, climatic change is shifting geographic centers of species distributions (Hovick 2016) and native and agricultural plant phenologies (Richardson et al. 2013). Furthermore, woody plant encroachment causes regime changes from historical grassland regimes to shrubland or woodland regimes (Engle et al. 2008). Entire ecoregions in the southern Great Plains have changed to woodlands in the last century, and many ecoregions in the Great Plains are increasingly at the risk of fundamental ecological change and regime shifts in the future (Twidwell et al. 2013, Wonkka et al. 2019).

Ecologists have had a long-lasting interest in the relationship between disturbances and the responses of ecological communities (Pickett and White 1985, Reynolds 1993, Brawn et al. 2001). Efforts over the last decades have focused on how the structure and function of species assemblages confer stability to disturbances at local and regional scales (Donohue et al. 2013, Fried-Petersen et al. 2020). This research is more recently extended to explore community-disturbance relationships from a resilience perspective. Specifically, the cross-scale resilience model (Peterson et al. 1998, Sundstrom et al. 2018) is used to assess the influence of species structural and functional diversity on resilience for ecosystems (Allen et al. 2005, Angeler et al. 2015a) and landscapes (Angeler et al. 2015b, Roberts et al. 2019). A major tenet of the cross-scale resilience model is that it accounts for the hierarchical organization of ecosystems and therefore allows assessment of disturbance effects on resilience within and across scales of time and space in the system. Accounting for scale in the analysis of disturbance effects is of particular interest because disturbances manifest distinctly at different scales in the system (Nash et al. 2014). For instance, a hail shower may have significantly stronger impact on seedlings compared to trees in a forest (Angeler et al. 2018). Furthermore, disturbances can surpass critical thresholds, which, in addition to scale, is a fundamental component of ecological resilience (Baho et al. 2017), causing the system to shift form one attractor domain to another (i.e. regime shift). It is long recognized that regime shifts produce a fundamental reorganization of ecosystems (Holling 1973), manifested in distinctly different pattern-process relationships and feedbacks between regimes (Allen et al. 2014). These fundamentally altered structures and functions are evident, for instance, in altered abiotic conditions and community structure (Angeler et al. 2015c) and the scaling patterns present in the system (Spanbauer et al. 2014).

Much research has so far emphasized the role of dominant species in driving biodiversity and community dynamics in response to environmental change, assuming implicitly that they are key players in ecological dynamics while de-emphasizing rare species. In fact, some modeling approaches routinely used by ecologists exclude rare species to not distort ordinations (e.g., correspondence analysis) or disregard species that are not significantly related to model outcomes (e.g., correlated with canonical axes in redundancy analysis) (but see Baker and King 2010). However, there is mounting evidence that rare species can play an important role in maintaining ecological pattern-process relationships and thus adaptive capacity (i.e. the latent potential of an ecosystem to alter resilience in response to change) after disturbances (Angeler et al. 2019). Mouillot et al. (2013) found that rare species in alpine meadows, coral reefs, and tropical forests comprised functional trait combinations that were not represented by abundant species. These authors suggested that if rare species go extinct, negative effects on ecosystem processes might result from the subsequent loss of adaptive capacity. Such negative effects may occur even if biodiversity associated with abundant species is high (Mouillot et al. 2013). The importance of rare species is also evident in their ability to replace dominant species after perturbation and maintain ecological functions in the system, which in turn contributes to ecological resilience (Walker et al. 1999). For instance, rare shrub species with larger root crowns than dominant species were able to compensate for the loss of dominant shrub species to mechanical disturbance by re-sprouting prolifically, thus maintaining a shrub-dominated system despite disturbance (Wonkka et al. 2016). This example shows that rare species may provide a relevant but, to some extent, unpredictable degree of an ecosystem’s capacity to adapt to change.

In this paper, we study the relevance of rare species in the context of spatial regime shifts associated with recently demonstrated shifting biome frontiers in the Great Plains of America caused by global change (Roberts et al. 2019). Because ecological systems undergo profound reorganization with persistent changes in structure, functions and feedbacks (Angeler and Allen 2016), we were especially interested in assessing whether rare species of one spatial regime might become dominant once a system has shifted to a new spatial regime, as is the case when grasslands become encroached by woodlands. We also determine whether rare species from one regime remain rare after the regime shift. Understanding these patterns may provide better insight into potential ecological legacies (Johnstone et al. 2016) that rare species from an old regime might leave in a new regime, specifically how they affect critical elements of adaptive capacity and resilience in the novel regime. To address our research question, we use time series modeling that infers the temporal scaling structure, and thus the hierarchical patterns necessary for assessing cross-scale aspects of resilience, and the dominant taxa that are contributing to these scale-specific dynamics (“deterministic species”) (Angeler et al. 2009). The modeling also allows for the identification of rare species which, because of their stochastic temporal dynamics (“stochastic species”), are unrelated to any scaling pattern identified (Baho et al. 2014). These stochastic species are considered to encompass rare taxa in this study, where rarity is defined as species occurring along a gradient from frequent incidences with low abundances to sporadic occurrences with higher abundances throughout the study period. That is, we consider time series modeling as an objective approach to differentiate between rarity and dominance based on a time-explicit criterion (i.e. over a defined study period; see below).

We study the relevance of potential ecological legacies of stochastic species using breeding bird communities in the North American Great Plains as a model system. Because of the south-to-north movement of ecoregions or spatial regimes (Roberts et al. 2019), we assess how stochastic species of the northern regime may potentially influence stochastic and deterministic species patterns once it gets encroached by the expanding southern regime, turning into a novel spatial regime. We used a time-for-space substitution, an approach commonly used in ecology (Pickett 1989), especially in a climate change context (Blois et al. 2013). Modern regime shifts often unfold at time scales that are not covered by routine monitoring (Spanbauer et al. 2014). Space-for-time substitutions overcome this common problem in regime shift research by comparing spatially independent units that already occur in alternative regimes. In our specific case, as the southern regime is suggested to eventually expand into the northern regime with ongoing climate change during the next decades (Roberts et al. 2019), we assess if and how many stochastic species of the “vulnerable” (northern) regime will show either stochastic or deterministic patterns in the “expanding” (southern) regime. We also assess if these species occur at one or different temporal scales of the expanding regime, which allows us to determine their contributions to cross-scale resilience in the expanding regime. This might provide insight into the dynamically changing spatial resilience of landscapes and how this might affect management and conservation efforts for ecosystems and species with ongoing climate change (Cumming 2011, Allen et al. 2016). We also assess the conservation status of species contributing to the observed patterns to provide information about species conservation implications associated with spatial regime changes.

## Material and methods

### Data and study setup

We collected 47 years of data ranging 1968 - 2014 from the U.S. Geological Survey Breeding Bird Survey data (BBS) of North America. The data contain avian community composition that is collected by qualified observers along georeferenced, permanent roadside routes across North America (Sauer et al. 2017). These data are publicly available. Along each *ca* 39.5 km route, observers make 50 stops once every 0.8 km where they conduct point-count surveys. During each survey, observers stand at the stop and record for three minutes the abundance of all bird species that are visually or aurally detected within a 0.4 km radius. Surveys start thirty minutes before local sunrise and last until the entire route is finished. To increase uniformity in probability of bird detection, surveys are conducted only on days with little or no rain, high visibility, and low wind.

We removed all aquatic species from the Anseriformes, Gaviiformes, Gruiformes, Pelecaniformes, Phaethontiformes, Phoenicopteriformes, Podicipediformes, Procellariiformes, and Suliformes families from the analysis because of known negative observation biases for waterfowl compared with terrestrial avian families (Holling 1992). We also removed hybrids and unknowns, and we condensed subspecies to their respective species.

Time series data across routes were heterogeneous with many missing data, but three transects were found suitable for analysis for each of the two regimes studied. The three transects, which together comprised 82% of the total species pool in the southern regime and 86% in the northern regime, were averaged to obtain exhaustive species occurrences and to facilitate the comparison of species distributions between spatial regimes. For this study, we selected a southern (latitudes: 28.9 – 29.7; Western Gulf Coastal Plains) and a northern spatial regime (latitudes: 31.8 – 33.4; South Central Plains) for analyses (Figure 1). These regimes were chosen because of documented biogeographical shifts (northward movement of southern regimes) with ongoing climate change (Roberts et al. 2019).

**Figure 1:**
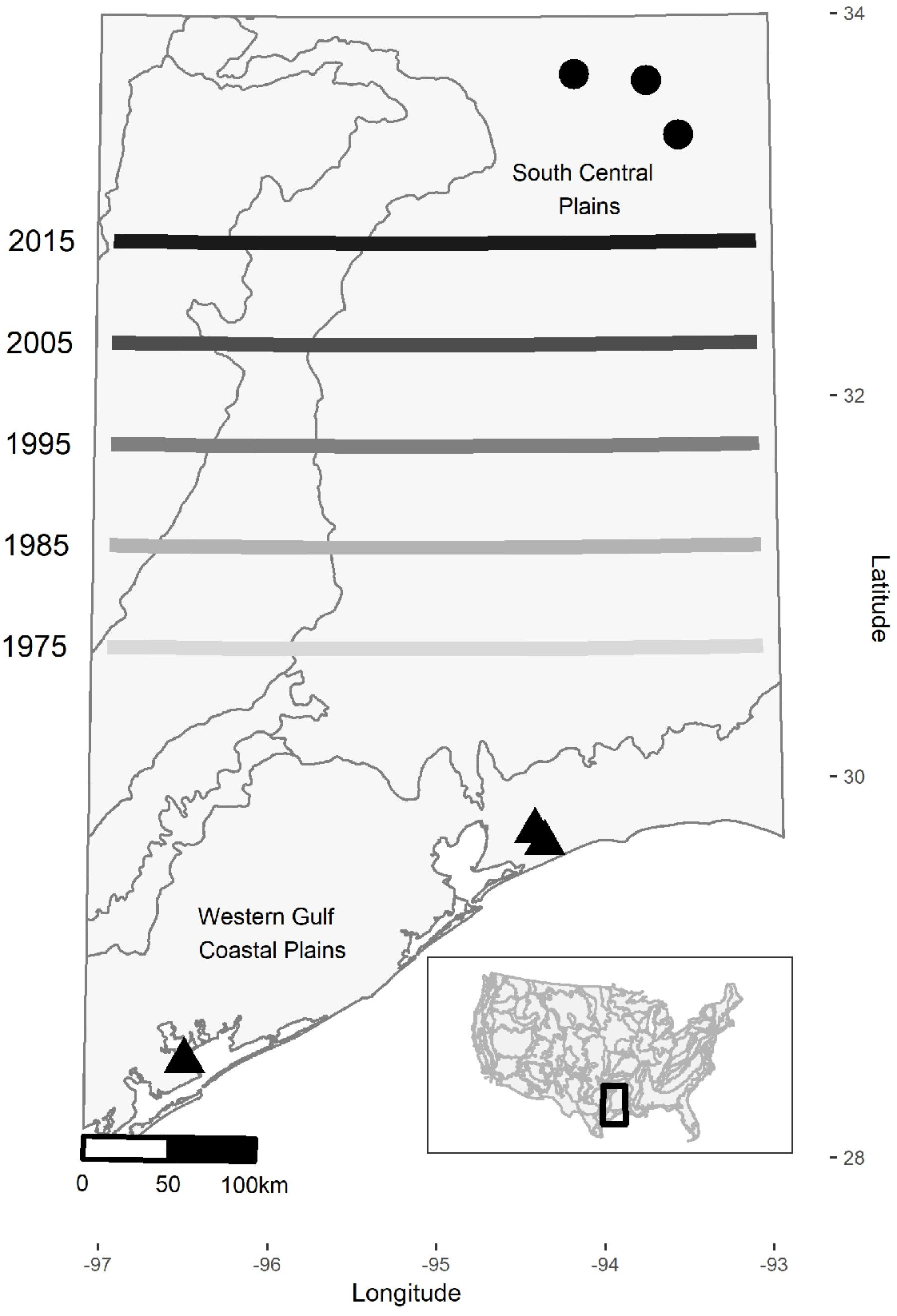
Map showing location of breeding bird survey transects in two spatial regimes. Contours show level 3 eco-region delineations based on US EPA and lines denote northward movements of the southern regime front over time assessed by Roberts et al. (2019).

### Statistical Analysis

#### Time series models

All statistical analyses were carried out in R 3.0.2 (R Development Core Team 2012) using the ‘aem.time’ function (AEM package, Blanchet and Legendre (2013)), packages nlme (Pinheiro et al. 2008), and the ‘quick PCNM’ function (PCNM package, Legendre et al. 2013). Asymmetric Eigenvector Maps (AEM) were extracted from a set of orthogonal temporal variables that were calculated from the time vector consisting of 47 steps between years 1968 and 2014. These AEMs are used as explanatory variables to model temporal relationships in the BBS data. In the case of AEM, the first variable models linear trends and subsequent variable show sine-wave patterns (Legendre and Legendre 2012), which allows assessing directional change and different inter-annual and decadal variation in the BBS. These extracted temporal variables are then used as explanatory variables in the time series models using redundancy analysis (RDA) (Angeler et al. 2009). Two time series models were constructed, one for the southern regime and one for the northern regime, which consisted of 122 species and 111 species, respectively, as response variables.

RDA selects significant temporal variables (AEMs) using forward selection. The selected variables are linearly combined in the RDA models to extract temporal structures from the bird species matrices. The modeled temporal patterns that are extracted from the data are collapsed onto significant RDA axes, which are tested through permutation tests. These RDA axes are then used to distinguish deterministic from stochastic species in the analysis. The R software generates linear combination (lc) score plots, which visually present the modeled temporal patterns that are associated with each RDA axis. That is, individual RDA axes indicate fluctuation patters at different temporal frequencies or scales. All bird species raw-abundances averaged from three transects per regime were Hellinger transformed prior to the analysis (Legendre and Gallagher 2001).

#### Correlation of Bird Taxa with Modeled Spatial Patterns

Following Angeler et al. (2015), we used Spearman rank correlation analysis to relate the raw abundances of individual bird taxa with the modeled temporal patterns (lc scores) associated with the RDA axes of both models. In this way we identified taxa that contributed significantly to the temporal dynamics revealed by the RDA (that is, deterministic species). Those taxa that were not associated with any significant canonical axis were identified as stochastic species.

Next, we examined whether stochastic species from the northern regime occur as either deterministic and/or stochastic patterns of the southern regime. This provides insight regarding how species change may alter the resilience of the northern regime once it is invaded by the southern regime with climate change. We also assessed the conservation status of these species according to the IUCN Red List (https://www.iucnredlist.org/) and the US Endangered Species Act (https://www.fws.gov/endangered/species/).

## Results

The analyses from the time series modeling of BBS data revealed significant temporal structure in both spatial regimes between 1968 and 2014. The overall variance explained by AEM models was high (adjusted R^2^ values; northern regime, 0.65; southern regime 0.63). The models revealed fluctuation patterns at distinct temporal scales (Figure 2). AEM revealed temporal dynamics associated with five and six significant RDA axes for the northern and southern spatial regime, respectively. Comparing both regimes, the temporal patterns were similar at RDA 1 and RDA 2. RDA 1 displayed a marked component of monotonic change in community composition. RDA 2 showed hump-bell shaped patterns. The remaining RDA axes indicated higher temporal variability of bird community structure within and between both regimes (Figure 2).

**Figure 2:**
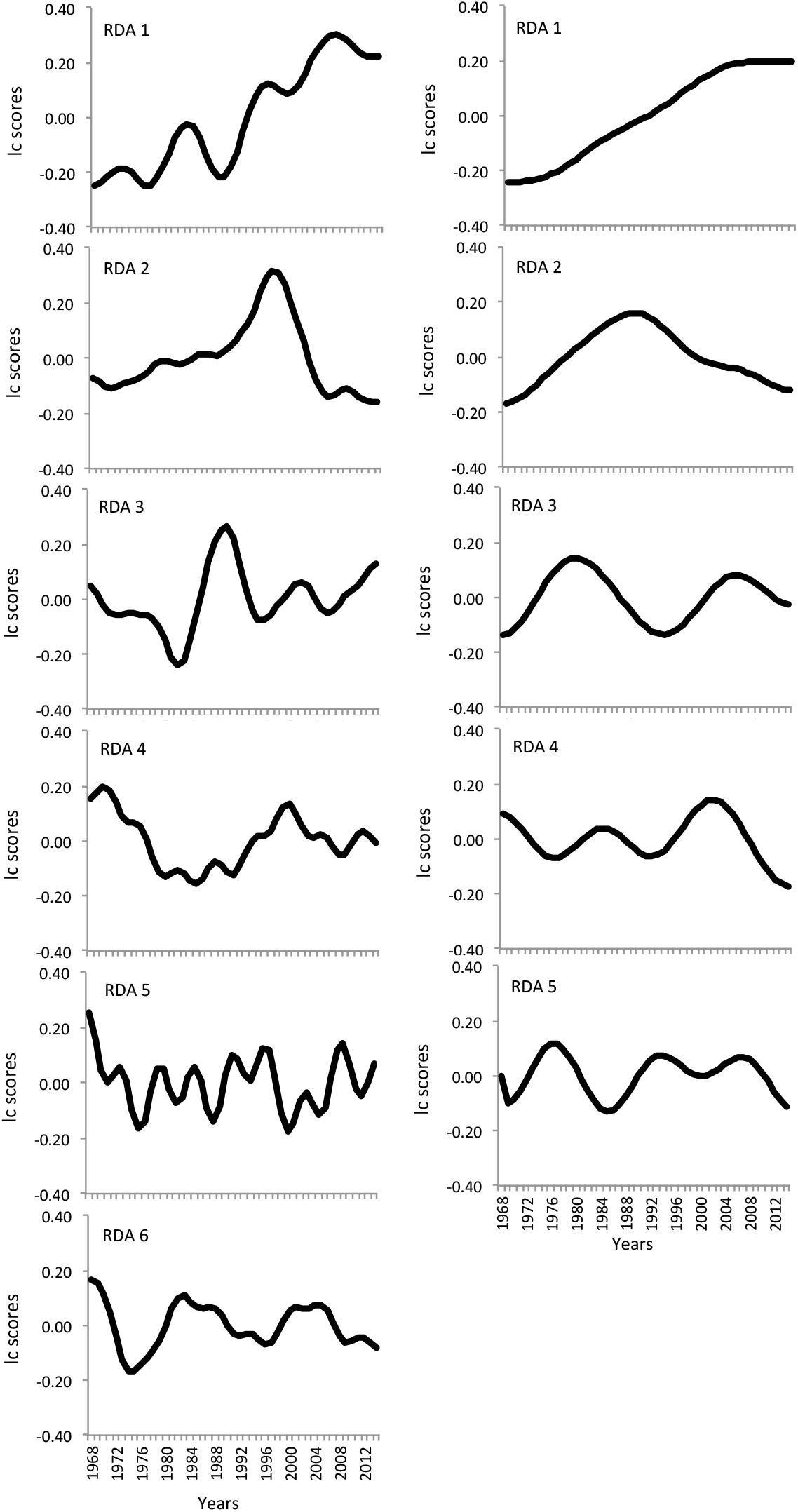
Linear combination (lc) score plots showing significant temporal patterns associated with different RDA axes in the time series models. Panels on the left represent the northern regime that becomes vulnerable to the invasion of the southern regime (right panels) due to climate change.

From the 122 bird species present in the southern regime, 87 (71%) were deterministic and 35 (29%) stochastic. From the 110 taxa present in the northern regime, 86 species (78%) were deterministic and 24 taxa (22%) stochastic. Across deterministic species, most taxa were correlated with RDA 1 (56 northern regime, 45 southern regime), followed by RDA 2 (19 northern, 12 southern) (Table 1). Only a few, generally fewer than 8, species correlated with the remaining RDA axes, except RDA 4 (19 species) of the southern regime model (Table 1).

**Table 1.** Number of bird species correlating significantly (P ≤ 0.05) with significant RDA axes in time series models (i.e. deterministic species) of the northern and southern regimes. Shown are also the number of stochastic species (uncorrelated with any RDA axis), and the total number of species in the data sets.

| Regime | RDA 1 | RDA 2 | RDA 3 | RDA 4 | RDA 5 | RDA 6 | Deterministic | Stochastic | Total |
| --- | --- | --- | --- | --- | --- | --- | --- | --- | --- |
| Northern | 56 | 19 | 4 | 4 | 3 | -- | 86 | 24 | 110 |
| Southern | 45 | 12 | 8 | 19 | 1 | 7 | 87 | 35 | 122 |

From the 24 stochastic species present in the northern regime, 10 were also found in the southern regime (Table 2), meaning that less than half of the stochastic taxa of the former regime occurred in the latter. From these, 5 species (*Thyromanes bewickii, Bubo virginianus, Pandion haliatus, Tachycineta bicolor*, *Vireo flavifrons*) remained stochastic and 5 species correlated with different RDA axis of the southern regime (Table 2). Specifically, *Dumetella carolinensis* and *Buteo swaisoni* correlated with RDA 1, *Emerophia alpestris* and *Passerina ciris* with RDA 2, and *Sethophaga americana* with RDA 3 while no stochastic species of the northern regime were found at RDAs 4, 5 and 6 of the southern regime model. This indicates that stochastic species can become deterministic at different scales in a new regime. None of these species were found to be of conservation concern according to the IUCN red list and the Endangered Species Act.

**Table 2.**
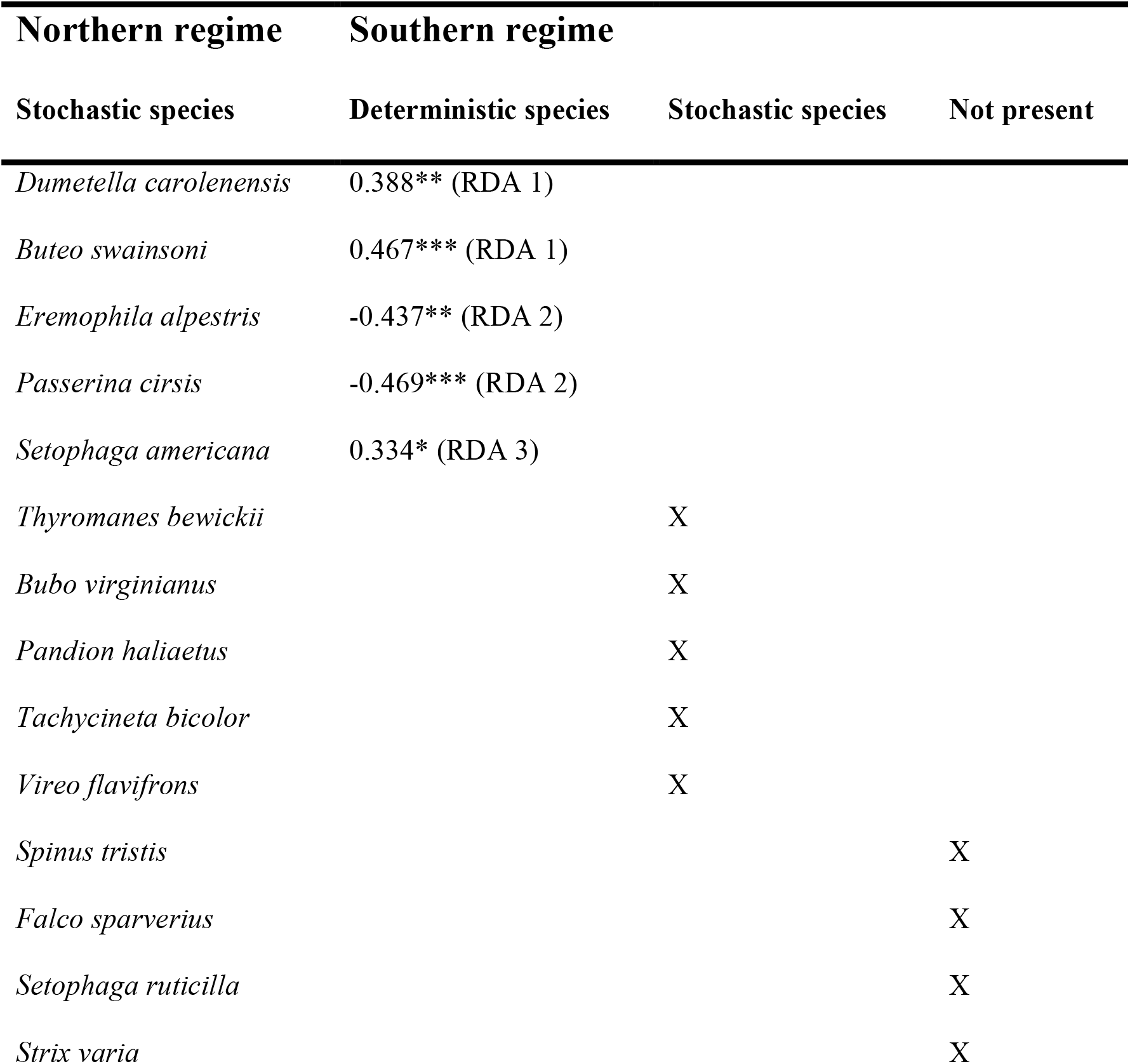
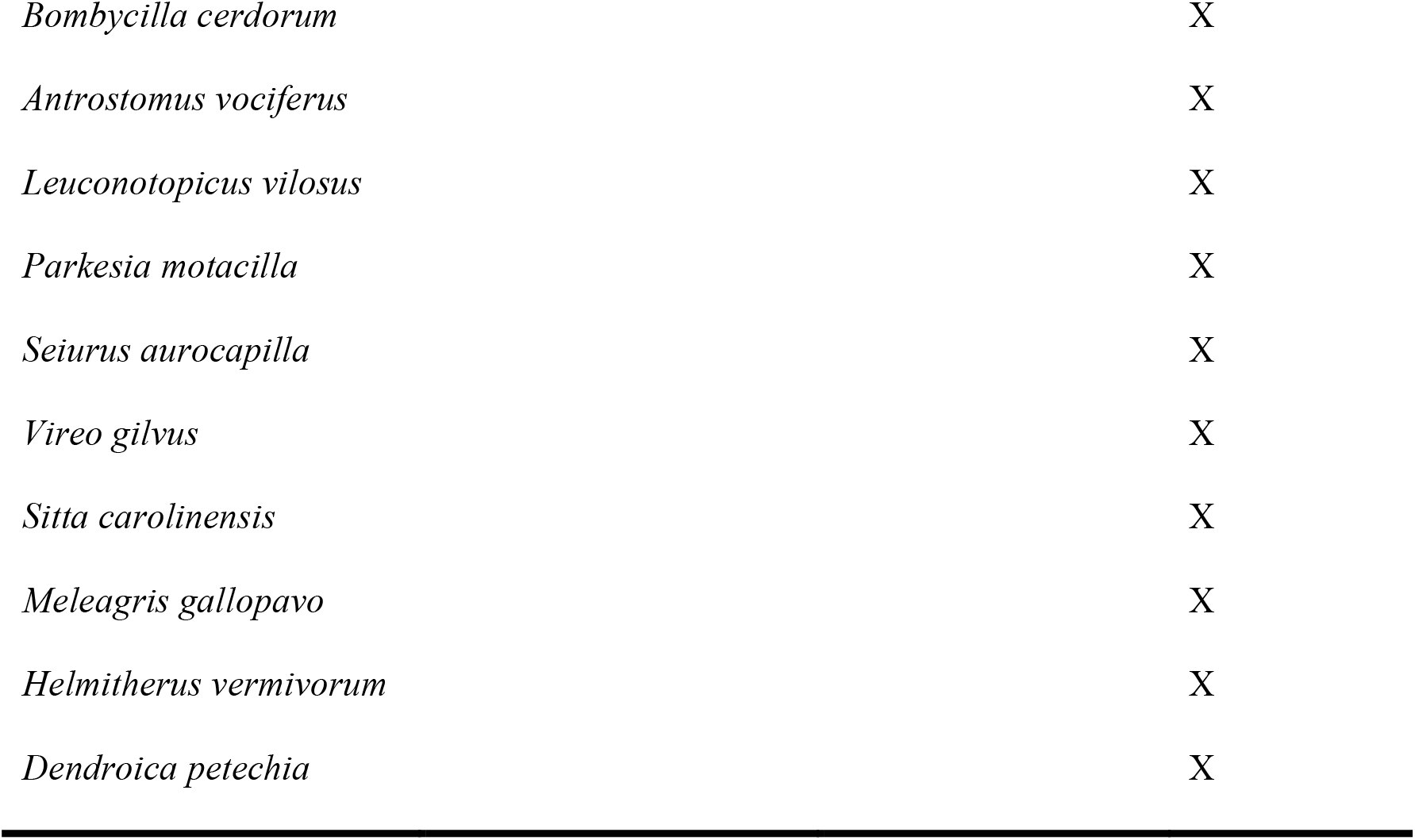
Stochastic species from the northern regime and their occurrences as deterministic or stochastic species in the southern regime revealed by Spearman rank correlation analysis. For deterministic species, Spearman Rho correlation coefficients, P levels (* P ≤ 0.05, ** P ≤ 0.01, *** P ≤ 0.001) and associated RDA axes from the time series model are shown. Northern stochastic species that remained stochastic (no significant correlations with RDA axes) or were not present in the southern regime are also shown (indicated with X).

## Discussion

That ecological systems undergo substantial structural and functional change after regime shifts is increasingly supported by empirical evidence (e.g., Spanbauer et al. 2016). The results of this study build on this existing body, adding an ecological legacy component (Johnstone et al. 2016); specifically, how stochastic species from one spatial regime might contribute to the adaptive capacity and resilience of a new spatial regime.

We use time series modeling to quantify resilience in both spatial regimes based on the cross-scale resilience model that helped us distinguish from a time explicit point of view between abundant and rare species through an objective, model-based distinction of deterministic and stochastic species in the data set (Baho et al. 2014). Furthermore, the approach allowed us to assess temporal scales present in the regimes and at which scales individual species fluctuate. This distinction is an objective representation of dynamics of abundant and rare species in the community (Baho et al. 2014), which allows for a refined assessment of community change opposed to approaches that are not time explicit and based on arbitrary delineations of species rarity or whole communities (Gaston 1994). Results show that deterministic species dominated the temporal dynamics in both spatial regimes, with the contribution of stochastic species being below 29%. While these results deviate from the notion that rare species dominate ecological communities (Magurran 2013), we emphasize that our results are based on modeling which explicitly accounts for species abundance changes and redundancies in these patterns over a defined time period. The approach therefore differs from methods based on snapshot samples that require different methods for assessing rarity.

Results revealed that only 10 out of the 24 stochastic species of the northern regime occurred in the southern regime. This finding is generally consistent with regime shift theory and further supports that regime shifts cause substantial ecological reorganization (Angeler and Allen 2016). A novel finding of our study is the partitioning of these species between deterministic and stochastic patterns in the new regime. The modeling revealed that 5 stochastic species from the northern regime were also stochastic and the other 5 deterministic in the southern regime. These 5 species were associated with three different temporal scales in the southern regime model. Because these scales were relatively species rich, our results suggest that these stochastic species from the northern regime contribute a relatively low degree of within-scale and cross-scale resilience and adaptive capacity in the southern regime. It is therefore reasonable to assume that such a pattern will be produced once the southern regime has invaded the northern regime. It has been suggested that rare species can compensate for the loss of dominant species and maintain the adaptive capacity of ecosystems after disturbances (Walker et al. 1999, Wonkka et al. 2016). Our results show that this does not necessarily need to be the case when systems undergo a regime change. Only 10 stochastic species of the northern regime were represented in the bird community of 122 species of the southern regime, which suggests that stochastic species from the northern regime may only leave a marginal ecological legacy once the northern regime becomes invaded by the southern regime.

Our study is based on space-for-time substitutions in which a regime change is only implicit. That is, rather than explicitly assessing regime changes resulting from northward migration of spatial regimes, our approach compared community dynamics between regimes, assuming that over time the northern regime will be encroached by the southern regime. Given our study design, the potential role of ecological legacies is therefore also only indirectly assessed because we lack the explicit sequential replacement of regimes and their species pools. While space-for-time substitutions have been criticized, they are still a valuable alternative to long-term studies (Pickett 1989), particularly in a climate change context (Blois et al. 2013). Space-for-time substitutions are therefore particularly useful for regime shift research, which is often limited by monitoring data that do not cover relevant scales of ecological change, which can be slow, particularly in a spatial context. There is need to account for both space and time in the assessment of spatial resilience (Cumming 2011, Allen et al. 2016, Sundstrom et al. 2017). This further underscores the utility of space-for-time substitutions as an important approach in spatial regime shift research, especially given the fast biogeographical changes on a rapidly changing planet.

We conclude by highlighting that none of the stochastic bird species of the northern regime contributing to deterministic or stochastic patterns in the southern regime were of conservation concern according to the ICUN Red list and the United States Endangered Species Act. From the perspective of conservation biogeography, this suggests that likely no urgent adaptations of species conservation plans are needed once the northern regime becomes invaded by the southern regime. However, careful interpretation of this implication is needed because our study design and results are agnostic to the potential future biodiversity changes and threat status that may occur beyond the temporal horizon of our study.

## Acknowledgements

This work was supported by Department of Defense Strategic Environmental Research Development Program W912HQ-15-C-0018, Nebraska Game & Parks Commission W-125-R-1 and the University of Nebraska-Lincoln’s Institute of Agriculture and Natural Resources. Additional support was provided by a sabbatical professorship to DGA by the University of Nebraska – Lincoln.

